# Toward Accurate RNA Folding Thermodynamics: Evaluation of Enhanced Sampling Methods for Force Field Benchmarking

**DOI:** 10.64898/2026.01.19.700441

**Authors:** Petra Kührová, Vojtěch Mlýnský, Michal Otyepka, Jiří Šponer, Pavel Banáš

## Abstract

Biologically functional RNAs operate near marginal stability, and their rugged free-energy landscapes and profound structural dynamics – typically not captured by structural biology experiments – play decisive roles. Atomistic molecular dynamics (MD) simulations provide a unique means to characterize these features. However, the applicability of atomistic MD is currently limited by accessible simulation timescales and, most importantly, by force-field (FF) accuracy. Folding free energies (ΔG°_fold_) of small RNA motifs represent well-defined targets for quantitative benchmarking of RNA FFs. In practice, however, obtaining thermodynamic estimates that are sufficiently robust for direct comparison with experimental data remains highly challenging, even for small RNA systems, and many published studies rely on sampling that is not fully converged. Here, we systematically assess the performance of widely used advanced enhanced-sampling techniques using the 8-mer r(gcGAGAgc) tetraloop as a representative benchmark system. We test temperature replica exchange (T-REMD), two solute-tempering variants of replica exchange (REST2 and REHT), as well as well-tempered metadynamics and on-the-fly probability enhanced sampling combined with solute tempering (ST-MetaD and ST-OPES). Among the tested approaches, T-REMD proves to be the most robust, yielding reproducible folding equilibria and consistent estimates of ΔG°_fold_ after approximately 20 μs of simulation time, independent of the initial folded or unfolded conformational ensemble. Our results provide practical guidelines for selecting sampling protocols suitable for quantitative RNA benchmarks and lay the foundation for systematic validation and future refinement of RNA FFs.

## INTRODUCTION

RNA molecules form dynamic ensembles of interconverting structures and play essential roles in diverse biological processes across viruses and living organisms. Their functions are closely tied to the ability to fold into complex three-dimensional architectures, stabilized by a delicate balance of intra- and intermolecular interactions.^1–4^ Molecular dynamics (MD) simulations have become an indispensable complement to experimental techniques such as X-ray crystallography, or NMR and FRET spectroscopies, providing atomistic insight into RNA structural dynamics and frequently supplying the missing detail needed to interpret experimental observations.^5–10^ The reliability and predictive power of such simulations critically depend on the accuracy of the underlying force field (FF).^8, 11–13^ While MD simulations are routinely used to characterize local conformational dynamics, the development and validation of FFs require tackling far more demanding predictive folding simulations, which test not only the local structure and flexibility but also the global thermodynamic balance among structurally distinct states across the entire conformational space.^12, 14–27^ Despite extensive progress over the past three decades, no systematic and quantitative benchmarking framework has yet been established to validate RNA FFs and assess the reliability of simulation methods.

After parametrizing the first nucleic acid FF version for explicit solvent nucleic acids simulations,^28^ the development of nucleic acids FFs has historically focused on correcting clearly artificial behaviors observed in longer MD simulations.^6, 8, 10, 11^ For example, the bsc0 correction eliminated spurious irreversible α/γ backbone flips in B-DNA helices,^29^ while the OL3 reparameterization of χ dihedrals^30^ prevented the irreversible transition of A-RNA stems into ladder-like structures.^31, 32^ Current state-of-the-art RNA FFs, however, are limited not by such obvious pathologies in canonical helices but by numerous more subtle imbalances in non-bonded interactions that determine the thermodynamic stability and rugged free-energy landscape of diverse RNA motifs. These include for example (i) overstabilization of sugar–phosphate interactions,^17, 20^ (ii) insufficient stabilization of base pairing,^16, 19–21, 23^ (iii) over-repulsion of a weak –CH…O– interactions,^25, 33, 34^ and/or (iv) imprecise electrostatics affecting ion-binding properties.^21, 35, 36^ A major issue is that manifestation of these imbalances is often non-uniform across diverse RNA systems and their impacts may be interdependent.^8, 12, 33, 37^ Some targeted corrections have been proposed and robustly tested. They include, for example, a modification of van der Waals parameters of phosphate oxygen atoms (the CP adjustment)^38^ reducing overstabilized phosphate electrostatic interactions^39, 40^ and gHBfix potentials introducing additional hydrogen-bond terms to address the underestimated base-pairing strength and excessive sugar–phosphate attraction.^20, 23^ Latest attempts implementing FF corrections to the standard OL3 RNA FF benefited from machine learning approaches and/or direct inclusion of experimental data into fitting schemes.^23, 27^ While earlier corrections could be validated by assessing local structural dynamics around native conformations, modern FF development increasingly requires accurate evaluation of the thermodynamic balance between folded, misfolded, and unfolded ensembles across the RNA conformational space.^8, 12, 22^

To advance this next stage of development, systematic benchmarking is essential. In analogy to popular benchmark sets commonly used in quantum chemistry,^41–44^ we envision the creation of an MD RNA benchmark set spanning a representative range of structural contexts. Such a set should include A-RNA duplexes as canonical references, small oligonucleotides to probe backbone flexibility, stable hairpin and internal loops such as tetraloops or the sarcin-ricin loop with its S-turn and GpU platform, flexible non-canonical motifs like kink-turns, and tertiary interaction motifs such as kissing loops or tetraloop–tetraloop receptor interactions.^12, 13, 45^ Crucially, each system must meet three criteria: (i) experimental characterization with sufficiently accurate thermodynamic data, ideally through UV-melting or extensive NMR data; (ii) the ability of simulations to reach converged predictions of these observables; and (iii) conditions under which the FF remains the sole determinant of agreement with experiment. Only then can benchmarking be objective and reproducible, providing a robust foundation for the next generation of RNA FF development.

Among the candidate systems, GNRA (Figure 1) and UNCG tetraloops (TLs) are among the most frequently used motifs in RNA FF testing.^14–16, 18–22, 25, 34, 46–48^ Their thermodynamic parameters, such as melting temperature and folding free energy (ΔG°_fold_), are either directly available from experiments^49–53^ or can be derived from Turner’s thermodynamics parameters.^54–57^ TLs are simple non-trivial RNA motifs of reasonably small size that combine canonical and non-canonical interactions, extending beyond the simplest oligonucleotide and duplex tests. These features make them highly attractive candidates for a benchmark set. To serve this role, simulations must be able to quantitatively predict their thermodynamic stability within a given FF. However, this goal has proven difficult to achieve, as previous enhanced-sampling studies of TLs have consistently suffered from incomplete convergence, preventing reliable estimation of ΔG°_fold_ and melting temperatures.^15–17, 20, 22^ Standard MD simulations are limited by accessible timescales and therefore often fail to sample rare events and slow conformational transitions. Enhanced sampling methods address this limitation by promoting more efficient exploration of configurational space and can be broadly classified into two main categories: (i) annealing replica-exchange methods and (ii) importance sampling approaches. Annealing replica-exchange methods accelerate sampling by increasing the real or effective temperature, thereby facilitating enthalpy barrier crossing without requiring prior knowledge of slow degrees of freedom. Frequently applied examples include temperature replica exchange MD (T-REMD)^60^ and Hamiltonian-based variants such as replica exchange with solute tempering (REST2)^61^, which selectively scale solute interactions. Related approaches, including accelerated or flooding-based methods (e.g., accelerated MD; aMD^62^, Gaussian-accelerated MD; GaMD,^63–65^ etc.), modify the potential energy landscape to mimic temperature effects. Annealing replica-exchange approaches are robust and broadly applicable, particularly for enthalpy-dominated processes, but are less effective for entropy-driven transitions and typically require careful parameterization (e.g., temperature ladders or scaling factors), often at substantial computational expense. Importance sampling methods enhance sampling by biasing a small set of carefully chosen collective variables (CVs) that describe the slow degrees of freedom governing rare events. Their efficiency critically depends on the quality of the selected CVs, requiring prior physical insight.^66, 67^ Importance sampling techniques include multi-window approaches such as umbrella sampling^68^ (usually combined with weighted histogram analysis method^69^), as well as single-trajectory methods like (well-tempered) metadynamics,^70, 71^ which constructs a history-dependent bias to flatten the free-energy landscape. While importance sampling methods can efficiently sample entropy-dominated processes and provide direct access to free-energy surfaces, missing or poorly chosen CVs can severely distort their performance. In addition, free-energy landscapes of biomolecules tend to be intrinsically multi-dimensional and their description is not always amenable to dimensionality reduction to one or two CVs. Interested readers are referred to recent reviews for detailed descriptions and practical applications of enhanced sampling methods.^8, 72–78^

**Figure 1.**
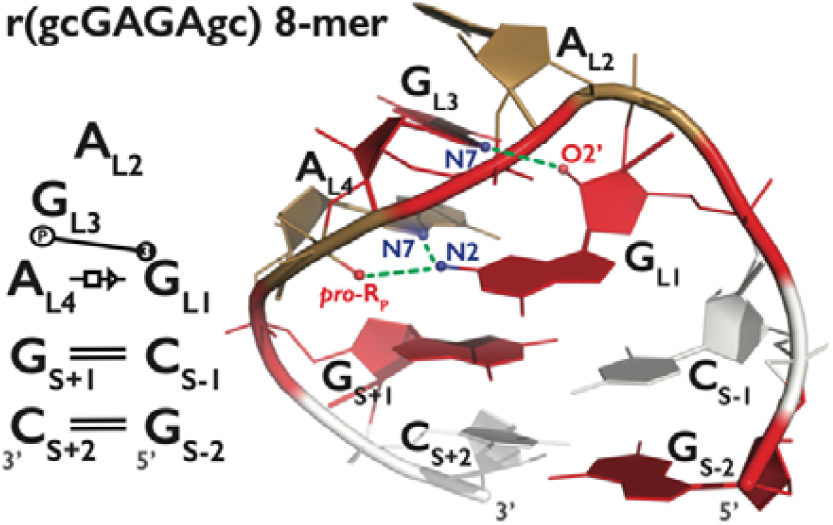
Secondary and tertiary structure of the 8-mer GAGA TL. Left panel presents secondary structure annotation according to the Leontis−Westhof nomenclature.^58^ It contains two canonical GC base pairs, the trans Hoogsten/Sugar-Edge (tHS)^58^ A_L4_/G_L1_ pattern, a purine triple base stack formed by A_L2_, G_L3_, and A_L4_, and the base-phosphate interaction type 3 (3BPh).^59^ Right panel shows the reference native state. A, C, and G nucleotides are colored in sand, white, and red, respectively. Signature H-bonds are colored in green.

In this work, we systematically evaluate several widely used enhanced sampling techniques for the folding of the 8-mer GAGA TL (Figure 1), assessing their ability to deliver converged and quantitatively robust thermodynamic descriptions. Our results highlight simulation strategies that enable incorporations of TL motifs into a future RNA benchmark set, helping bridge the gap between experimental data and predictive FF testing.

## METHODS

### Starting structure and simulation setup

The unfolded starting structures of the r(gcGAGAgc) 8-mer TL (GAGA TL) were taken from our previous REST2 simulations.^20^ Reference native state (i.e., the folded structure) was taken from the 1.04 Å resolution X-ray structure of the sarcin-ricin loop (PDB ID 1Q9A,^79^ residues 2658-2663) and capped by an additional one GC base-pair yielding the final r(gcGAGAgc) sequence. The starting structures were solvated using a rectangular box of OPC^80^ water molecules with a minimum distance between box walls and solute of ~10 Å. The final simulation boxes of both folded and unfolded systems were designed to have comparable size to ensure consistency in the analysis. Simulations were conducted at ~1 M KCl salt excess using Joung&Cheatham^81^ ion parameters for TIP4PEW water (K^+^: *R* = 1.590 Å, ε = 0.2795 kcal/mol, Cl^−^: *R* = 2.760 Å, ε = 0.0117 kcal/mol). We used hydrogen mass repartitioning which enables a 4-fs integration time step.^82^

The *ff*99bsc0χ_OL3_ (i.e., OL3)^28–30, 83^ RNA FF was employed and further adjusted by the van der Waals (vdW) modification of phosphate oxygens developed by Steinbrecher et al.^38^ in combination with our reparameterization of the affected α, γ, δ and ζ backbone torsions.^39^ This RNA FF version is further abbreviated as OL3_CP_ and corresponding AMBER library files are provided in the Supporting Information of Ref. ^16^. All simulations incorporated gHBfix19 potential,^20^ where all −NH…N– base– base interactions are strengthened by 1.0 kcal/mol and all −OH…bO– and −OH…nbO– sugar– phosphate interactions are weakened by 0.5 kcal/mol. We chose the original gHBfix19 correction over the more recent and computationally more demanding gHBfix21 potential,^23^ as prior works demonstrated that gHBfix19 is sufficient to reproduce the folding of the GAGA TL.^20, 22^

Among the tested enhanced sampling methods, T-REMD^60^ and REST2^61^ simulations were performed using AMBER20^84^ with the pmemd.cuda engine.^85^ Both folded and unfolded structures were prepared using the tLEaP module of the AMBER18^86^ program package. Simulations were performed in cubic box and the NVT ensemble (constant volume), with long-range electrostatics calculated using the particle mesh Ewald method,^87^ employing a 1 Å grid spacing and a 10 Å real-space cutoff. Exchange attempts were made every 10 ps, and Langevin dynamics with a friction coefficient of 2 ps^-1^ was used for temperature control. Hybrid replica exchange (REHT)^88^ and well-tempered metadynamics combined with REST2 (i.e., solute tempering with metadynamics, ST-MetaD)^22, 89^ were carried out using the GPU-enabled version of GROMACS2018^90^ in combination with PLUMED2.5,^91^ employing the Hamiltonian replica-exchange framework.^92^ The on-the-fly probability enhanced sampling (OPES)^93^ combined with REST2 (i.e., solute tempering with OPES, ST-OPES) required more recent PLUMED version and was therefore performed using GROMACS2022 in combination with PLUMED2.8. The simulation protocol in GROMACS differed slightly from that used in AMBER due to differences between the simulation engines. Specifically, GROMACS simulations were performed in a rhombic dodecahedral box, and all bonds involving hydrogen atoms were constrained using the LINCS algorithm.^94^ The cutoff distance for the real-space summation of electrostatic interactions was set to 10 Å, and temperature control was achieved using the stochastic velocity-rescaling thermostat.^95^

### Native state definition

The folded state was defined using an *ε*RMSD matric^96^ cut-off of 0.7 relative to the reference native structure. Structures with *ε*RMSD < 0.7 retain all native (signature) H-bonds, i.e., (i) the G_L1_(N2H)…A_L4_(*pro*-R_P_) base-phosphate interaction type 3 (3BPh),^59^ which can alternate with G_L1_(N1H/N2H)…A_L4_(*pro*-R_P_) bifurcated H-bonds (4BPh interaction), during the simulation, (ii) G_L1_(N2H)…A_L4_(N7) base-base, and (iii) G_L1_(2’-OH)…G_L3_(N7) sugar-base interaction. The A_L2_, G_L3_, and A_L4_ bases form a purine triple base stack and G_L1_ is base-paired with A_L4_ by the trans Hoogsten/Sugar-Edge (tHS)^58^ A_L4_/G_L1_ pattern (Figure 1).

### T-REMD Settings

We performed two T-REMD simulations, each employing 64 replicas, with one initiated from the native conformation and the other from unfolded (fully extended) conformations. The temperatures spanned the range from ~278 K to ~461 K and were chosen to maintain an exchange rate of ~25%. Both T-REMD simulations were run for 25 μs per replica.

### REST2 Settings

Three REST2 simulations with 16, 32, and 64 replicas were initiated from unfolded (fully extended) conformations. The scaling factor (λ) values ranged from 1.0908 to 0.658253, 0.6433105, and to 0.635839 for 12, 32, and 64 replicas, respectively. The corresponding effective temperature ranges were thus approximately from ~273 K to ~453 K, ~463 K, and to ~469 K for 12, 32, and 64 replicas, respectively. Each REST2 simulation was run for 20 μs per replica (Table 1).

**Table 1.** List of all performed enhanced sampling simulations.^a^.

| Method | Replicas | Time [μs] | Starting conformation |
| --- | --- | --- | --- |
| T-REMD | 64 | 25 | native |
|  | 64 | 25 | unfolded |
| REHT | 20 | 20 | unfolded |
| REST2 | 16 | 20 | unfolded |
|  | 32 | 20 | unfolded |
|  | 64 | 20 | unfolded |
| ST-MetaD | 12 | 4x5<sup>b</sup> | unfolded |
|  | 16 | 5 | unfolded |
|  | 12 | 2x30 | unfolded |
| ST-OPES | 12 | 30 | unfolded |

<sup>a</sup> All simulations were run with the OL3<sub>CP</sub> RNA FF and gHBfix19 potential. See Methods for settings of each enhanced sampling method and further details.
<sup>b</sup> Data from one (out of four) 5 μs-long ST-MetaD simulation with 12 replicas were taken from our previous work.<sup>22</sup>

### REHT Settings

One REHT simulation was performed starting from the unfolded conformation. In comparison with the REST2 approach, the REHT method differentially heats both the solute and the solvent, and was designed to reduce the residence times of metastable states in folding simulations of intrinsically disordered proteins.^88^ While the bath temperature is constant across all replicas in REST2 simulations, it is raised mildly up to ~340 K along the replica ladder in the REHT setup.^88^ Here we initially aimed to follow the REST2 setup with 16 replicas; however, the additional degrees of freedom arising from the explicit treatment of water in the REHT method affect the required number of replicas, which is moderately higher compared to the REST2 simulation. Therefore, we used 20 replicas with λ values ranging from 1.058401 to 0.599598, corresponding to an effective temperature range from ~282 K to ~497 K. REHT simulation was run for 20 μs per replica.

### ST-MetaD Settings

Seven ST-MetaD simulations with 12 or 16 replicas were initiated from unfolded conformations. The effective temperature range was set up from ~298 K to ~497 K and from ~273 K to ~453 K for 12 and 16 replicas, respectively. The *ε*RMSD metric was used as a biasing CV.^15, 22^ The augmented cutoff for biasing was set at 3.2 (*ε*RMSD_aug_) as this value was shown to allow forces to drive the systems towards and away from the native conformation.^15, 22^ *ε*RMSD with standard cut-off (2.4) was used for analysis, where snapshots with *ε*RMSD < 0.7 were considered as folded (native-like) ensemble. Sampling of each replica was enhanced by Gaussian depositing every 1 ps with a height ~0.5 kJ/mol and Gaussian width set to 0.1 (CV units). We rescaled the height of the Gaussians with a bias-factor of 15. Only the reference replica (with an effective temperature of 298 K) was used to estimate populations of the native state. ST-MetaD simulations were run for 5 μs or 30 μs per replica (Table 1).

### ST-OPES Settings

A single ST-OPES simulation employing 12 replicas was initiated from unfolded conformations. The effective temperature spanned approximately 298–497 K and the εRMSD_aug_ metric was used as the biasing CV, consistent with the settings adopted for the ST-MetaD simulations with 12 replicas. Additional simulation parameters closely followed those used for ST-MetaD. Sampling in each replica was enhanced by depositing bias every 1 ps, with the free-energy barrier parameter (BARRIER) set to 30 kJ/mol, a kernel width of 0.1 (in CV units), and a bias factor of 15. Analysis tools provided by the original authors^93^ were used to reconstruct the free-energy profile and to compute the folded-state population from the reweighted ensemble (https://github.com/invemichele/opes/tree/master/postprocessing). The ST-OPES simulation was run for 30 μs per replica (see Table 1 for a summary of all simulations performed).

## RESULTS & DISCUSSION

To assess the convergence and performance of enhanced sampling methods for RNA folding, we focus on the 8-mer GAGA TL as a representative benchmark system. We employ and compare five widely used enhanced sampling techniques, generating more than 7.2 ms of new simulation data. Our goal is to identify simulation strategies capable of delivering quantitatively converged and robust thermodynamic descriptions of this short RNA motif, thereby enabling its reliable use in future RNA FF benchmarking.

### Convergence of Annealing Replica-exchange Simulations

We tested three annealing replica-exchange approaches, i.e., T-REMD,^60^ REST2,^61^ and its derivative REHT,^88^ which are widely used to accelerate conformational sampling of RNAs. They enhance sampling by overcoming enthalpic barriers via elevated temperatures and/or Hamiltonian scaling, while maintaining a canonical ensemble at the reference replica. To assess their convergence, we performed extensive simulations of the GAGA TL and compared ensembles obtained from trajectories initiated in fully folded and/or fully unfolded states (Table 1).

Two independent 64-replica T-REMD simulations (25 μs per replica) were initiated from either all folded or all unfolded states. This design provides a stringent convergence test because only sufficiently long simulations should yield the same equilibrium ensemble. In both cases, the folded fraction at 298 K approached the same value after ~18 μs, indicating the convergence (Figure 2). The folded fraction over the final 7 μs (*ε*RMSD < 0.7; errors represent standard deviations) was 22 ± 2 % and 22 ± 3 % for the unfolded- and folded-start simulations, respectively (Figure 2). The corresponding free energies were 0.76 ± 0.07 kcal/mol and 0.74 ± 0.09 kcal/mol, respectively, in excellent agreement between the two different starting conditions (see Tables S1, S2 and Figure S1 in Supporting Information for detailed comparison of temperature-dependent populations of main conformational states between both independent T-REMD simulations). At equilibrium, the temperature ladder contained on average 6 to 7 folded replicas. Although exchange rates were high enough to redistribute states across replicas, individual folding and unfolding events occurred on the microsecond timescale, leading to slow convergence of the total number of folded replicas, and consequently of the folded fractions at the individual temperatures. After the convergence, fluctuations in the total number of folded replicas (5 to 8) persisted but produced only minor variations at particular temperatures. These results demonstrate that T-REMD can quantitatively converge the folding equilibrium of 8-mer GAGA TL, providing reliable ΔG°_fold_, although tens of microseconds per replica were required even with the sampling enhancement afforded by replica exchange.

**Figure 2.**
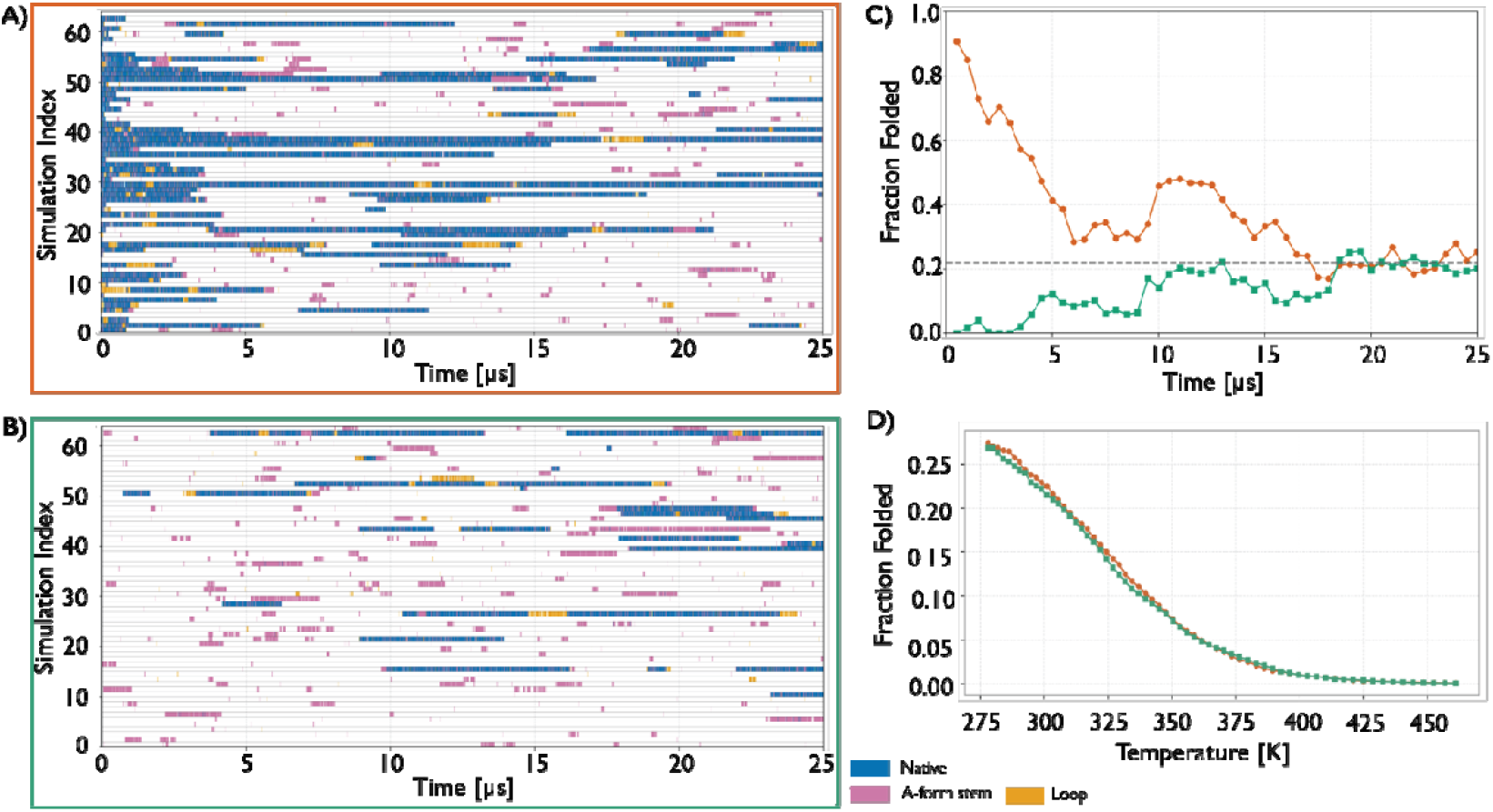
Convergence of T-REMD simulations of the GAGA TL. (A) Trajectories initiated from the folded state. (B) Trajectories initiated from unfolded states. In panels (A) and (B), each row corresponds to one of the 64 replicas, with structural states color-coded as follows: native state involving both stem and loop (blue), A-form stem (purple), and loop (orange). (C) Time evolution of the native-state population at 298 K for simulations initiated from folded (dark-orange) and unfolded (green) ensembles, evaluated at 0.5 μs intervals. The horizontal gray dashed line indicates the mean folded fraction of 22 %, averaged over the final 7 μs (εRMSD < 0.7). (D) Native state population as a function of temperature averaged over the final 7 μs.

REST2 simulations were performed with 16, 32, and 64 replicas (20 μs per replica) and were initiated from unfolded states. Overall, all REST2 simulations exhibited markedly slower and less reliable convergence than T-REMD. In the 16-replica case, the system entered an apparent steady regime after approximately 12 μs, with two folded replicas present in the ensemble. However, this regime cannot be considered converged, as evidenced by a significant decrease in the folded fraction during the final 2 μs of the simulation, when one folded replica unfolded (Figure 3A). The 16-replica ensemble contained only one to two fully folded replicas, which remained confined to the lower part of the temperature ladder and thus reduced the overall efficiency of the REST2 simulation.^22^ The small number of complete folding events amplified the influence of individual folding and unfolding transitions on the estimated folded fraction at 298 K, which did not fluctuate smoothly around a single value (Figure 3E). Instead, it exhibited pronounced jumps between discrete states defined by the number of folded replicas present in the temperature ladder. Averaged over the apparent steady-state regime corresponding to the final 8 μs of the simulation, the folded fraction was 20 ± 4 %, corresponding to ΔG°_fold_ of 0.82 ± 0.15 kcal/mol. Although the resulting standard error of the mean folded fraction is only moderately larger than that obtained from the T-REMD simulations, the presence of these discrete jumps highlights convergence issues and complicates the interpretation of the resulting thermodynamic estimates.

**Figure 3.**
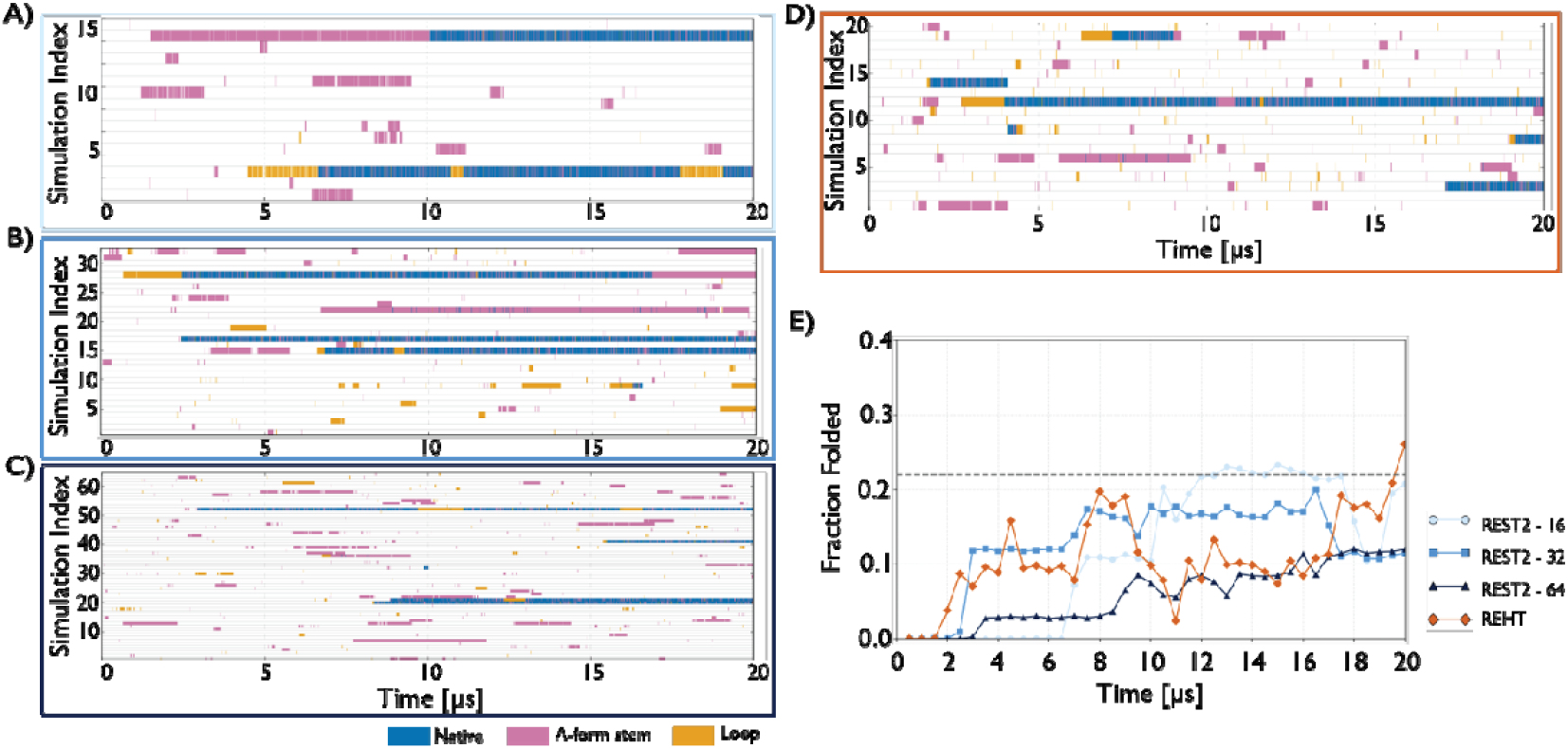
Convergence of REST2 and REHT simulations of the GAGA TL. (A–C) REST2 simulations employing 16, 32, and 64 replicas, respectively. (D) REHT simulation with 20 replicas. In panels (A–D), each row corresponds to one replica, with structural states color-coded as in Figure 2. (E) Time evolution of the native-state population at 298 K for the four simulations shown in panels (A–D): REST2 with 16 replicas (light blue), 32 replicas (blue), 64 replicas (dark blue), and REHT (dark-orange). The horizontal gray dashed line indicates the mean folded fraction of 22 % obtained from the T-REMD simulations.

Expanding the ladder to 32 replicas led to an apparent steady regime after approximately 7 μs, with two to three folded replicas present in the ensemble. The average folded fraction over the final 13 μs was 16 ± 3 %, corresponding to ΔG°_fold_ of 1.00 ± 0.12 kcal/mol. However, no substantial improvement relative to the 16-replica case was observed, as discrete jumps between different numbers of folded replicas and the associated fluctuations persisted (Figures 3B and 3E). As a result, these thermodynamic estimates remained less reliable than those obtained from T-REMD. By contrast, in the 64-replica simulation, the larger number of replicas substantially reduced these fluctuations, as approximately 6–7 folded replicas are expected at equilibrium, comparable to T-REMD. However, the approach to this regime was markedly slower (Figures 3C and 3E). Even after 20 μs, the folded fraction at 298 K continued to increase toward the T-REMD value without reaching a clear steady state. Thus, while increasing the number of replicas in REST2 simulations reduces statistical noise, it also delays convergence because a larger number of folding events across the temperature ladder is required to populate the higher equilibrium number of folded replicas. The slower kinetics observed in REST2 most likely arise from the selective scaling of solute interactions while the solvent remains unheated. This so-called “cold-solvent” effect slows solvent-mediated rearrangements and thereby hampers folding and unfolding transitions, even at the highest replicas.

To explicitly test whether tempering both solute and solvent can alleviate these limitations, we performed a 20-replica REHT simulation with 20 μs per replica. REHT combines temperature exchange with solute scaling, thereby enhancing also solvent sampling at higher replicas. After approximately 2 μs, the system appeared to enter a steady regime (Figure 3E), albeit with a folded fraction substantially lower than that observed in T-REMD. Only toward the end of the simulation, after about 17 μs, did the folded fraction begin to approach the T-REMD value. Throughout most of the trajectory, the ensemble contained only one folded replica (Figure 3D), and the folded fraction at 298 K exhibited pronounced jumps and large fluctuations due to the limited size of the replica ladder (Figure 3E). Averaged over the final 18 μs, the folded fraction was 12 ± 5 %, corresponding to ΔG°_fold_ of 1.17 ± 0.27 kcal/mol. Overall, while REHT partially mitigates the cold-solvent effect and accelerates kinetics relative to REST2, it remains strongly limited by the number of replicas and therefore does not provide the precision required for quantitative thermodynamic estimates.

Taken together, these results indicate that, among annealing replica-exchange approaches, only T-REMD achieves converged folding equilibria for the GAGA TL on accessible time scales (tens of microseconds per replica). In contrast, REST2 and REHT are limited either by restricted replica counts, which amplify statistical fluctuations, or by slow kinetics arising from incomplete solvent heating – or by both factors simultaneously. For short RNA hairpins, T-REMD therefore remains the most reliable annealing-based strategy for obtaining robust folding thermodynamics suitable for FF benchmarking, albeit at a substantial computational cost.

### Convergence of Importance Sampling Simulations

The second major class of enhanced sampling techniques relies on importance sampling along selected CVs. Both the employed ST-MetaD^22, 89^ and ST-OPES^93, 97, 98^ approaches combine solute tempering, which promotes extensive exploration of unfolded and misfolded states, with a bias applied to the εRMSD CV that quantifies the distance from the native structure. This combination ensures sufficient sampling of the folded state and unfolding/refolding events. The resulting simulations generate a non-canonical ensemble and a free-energy profile along the εRMSD CV. ΔG°_fold_ can be directly extracted from this profile, while reweighting procedures provide access to the unbiased canonical ensemble and thus to the populations of folded, misfolded, and unfolded states.

In contrast to annealing replica-exchange methods, both ST-MetaD and ST-OPES importance sampling approaches generate frequent transitions between folded and unfolded states due to the explicit bias applied along the εRMSD CV. As a result, convergence depends only weakly on the initial conformation, and all simulations were therefore initiated from unfolded structures. To assess convergence, we first performed five independent ST-MetaD simulations of 5 μs each (four using a 12-replica setup and one using 16 replicas). The folded fractions obtained from the reweighted ensembles of the reference replicas were 19 ± 16 %, 21 ± 15 %, 26 ± 26 %, 33 ± 21 %, and 31 ± 19 %. The reported standard deviations reflect variability within individual trajectories only, whereas the overall accuracy additionally depends on the convergence of the accumulated bias and the reconstructed free-energy profiles. Although the average folded fractions are broadly consistent across runs, the large statistical uncertainties, originating from pronounced fluctuations of the still-evolving bias potential, together with noticeable differences among the free-energy profiles, indicate that 5 μs-long sampling is insufficient to achieve quantitative convergence (Figure 4A, 4B). This limitation is particularly evident in the unfolded-state region, even when solute tempering is combined with MetaD to enhance exploration of misfolded and unfolded ensembles.

**Figure 4.**
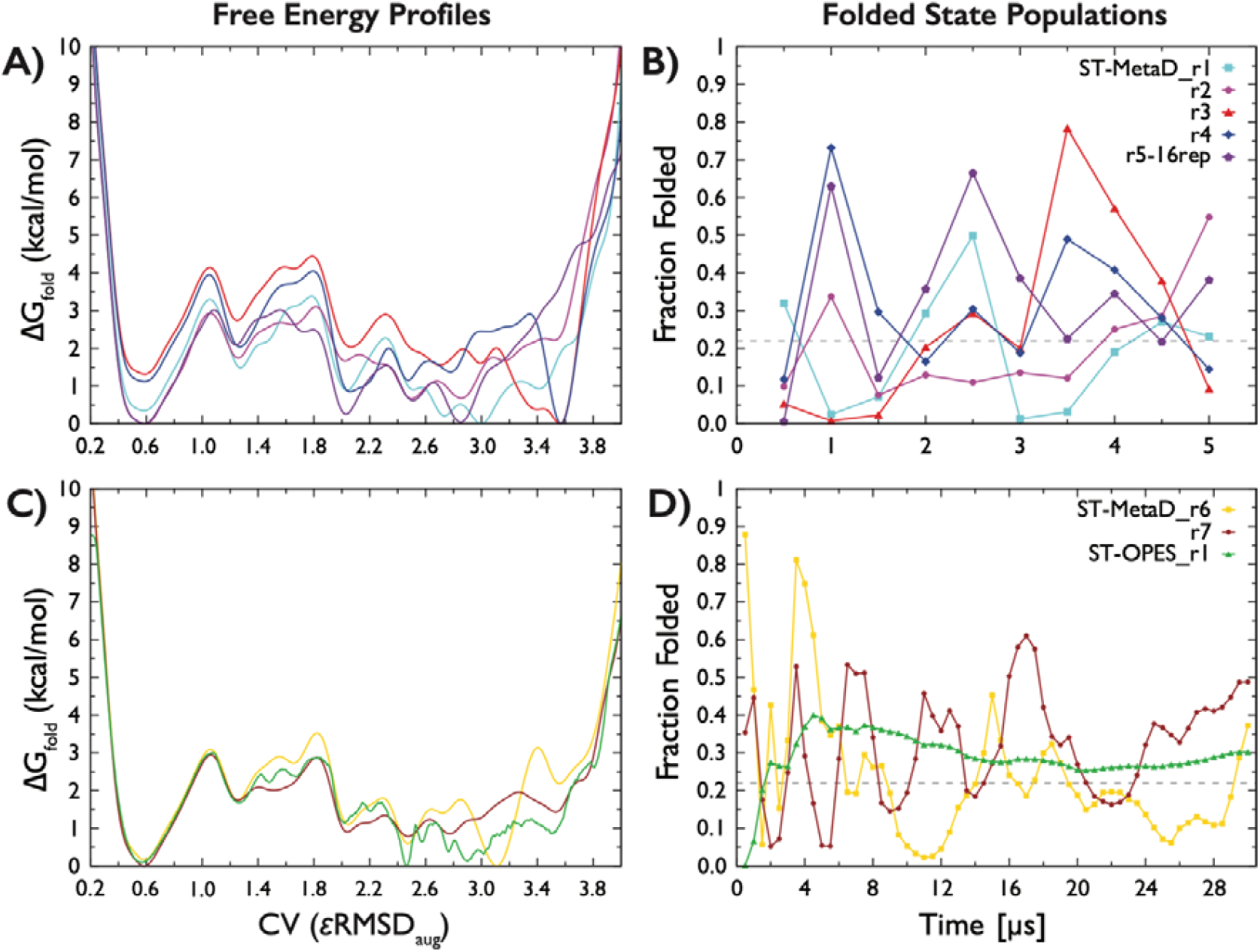
Convergence of ST-MetaD and ST-OPES simulations of the GAGA tetraloop. Free-energy profiles along the εRMSD_aug_ CV (see Methods) obtained from (A) five independent 5 μs-long ST-MetaD simulations and (C) two 30 μs-long ST-MetaD simulations and one 30 μs-long ST-OPES simulation, respectively. (B, D) Time evolution of folded-state populations estimated from the corresponding reweighted ensembles using standard εRMSD (see Methods), with each line representing values computed using the instantaneous bias accumulated up to the given time. Corresponding plots showing the time evolution of ΔG°_fold_ are provided in Figures S2 and S3 in the Supporting Information.

To improve convergence, we performed two additional simulations on longer time scales. Two independent 30 μs-long ST-MetaD trajectories yielded reweighted folded fractions of 24 ± 18 % and 32 ± 14 %, corresponding to ΔG°_fold_ of 0.8 ± 0.6 kcal/mol and 0.5 ± 0.5 kcal/mol, respectively. The convergence of the free energy profiles was substantially improved (Figure 4C), however, the error of the folding free energy and estimated folded state fraction remain high, indicating that full convergence has not yet been achieved. Compared with T-REMD, the ST-MetaD simulations still exhibit considerably larger statistical uncertainty in the folded-state populations and the corresponding ΔG°_fold_. In addition, we performed a 30 μs-long ST-OPES simulation, which yielded a folded fraction of 29 ± 6 %, corresponding to ΔG°_fold_ of 0.6 ± 0.5 kcal/mol. This result indicates that the ST-OPES method can mitigate fluctuations arising from free-energy minima overfilling in ST-MetaD and leads to improved convergence. This improvement is likely related to the fact that ST-OPES directly reconstructs free energies via Boltzmann reweighting of configurations sampled in the vicinity of each point on the CV surface.^93, 97, 98^ Nevertheless, in the case of the 8-mer GAGA TL, ST-OPES still performs worse than T-REMD simulations in terms of statistical uncertainty.

## CONCLUDING REMARKS

Although the GNRA TL represents one of the smallest and most frequently used RNA benchmark systems, achieving quantitative convergence of its folding equilibrium remains challenging.^14–16, 19, 21, 22, 27, 47, 48, 99–102^ In this work, we provide – for the first time – quantitative estimates of the ΔG°_fold_ of the 8-mer GAGA TL using the OL3_CP_–gHBfix19 FF, enabling direct comparison with experimental thermodynamic data. Among the tested enhanced sampling techniques, T-REMD emerged as the most reliable approach, yielding fully converged folding equilibria and quantitatively consistent thermodynamic parameters. In contrast, solute tempering methods, namely REST2 and REHT, were found to be less efficient. Their reduced replica ladder size, while constituting their main computational advantage, significantly hampers convergence. Slower intra-replica dynamics, most likely arising from the cold-solvent effect, further delay folding and unfolding transitions. As a result, complete folding–unfolding–folding transitions, which are required to achieve convergence,^22, 103, 104^ were never observed in continuous (demultiplexed) replicas on the tens-of-microseconds timescale (Figure 3). The combination of importance sampling methods with solute tempering, namely ST-MetaD and ST-OPES, drives the system close to equilibrium within a few microseconds, as the folded fraction rapidly approaches its expected equilibrium value. However, establishing a true equilibrium between folded and unfolded ensembles requires additional tens of microseconds, owing to slow convergence in the unfolded-state region (Figure 4). In the unfolded ensemble, multiple conformations overlap along the employed εRMSD CV and their mutual balance remain insufficiently sampled. While combining MetaD or OPES with solute tempering partially alleviates this limitation, it does not fully eliminate it. Consequently, under the present conditions, importance sampling techniques yield less precise thermodynamic estimates than T-REMD for the GAGA TL folding, despite being applicable with substantially fewer replicas and therefore a significantly lower total computational cost.

While annealing methods such as T-REMD proved most accurate for the GAGA TL, their efficiency strongly depends on the nature of the folding barrier. Optimal performance can be expected for processes dominated by enthalpic contributions and characterized by relatively low entropic barriers between folded and unfolded ensembles. In such cases – including the folding of small, monomeric RNA hairpins – T-REMD provides highly reliable estimates of folding thermodynamics. By contrast, when folding involves extensive conformational reorganization and significant entropic penalties, as in the formation of RNA duplexes, folding and unfolding kinetics are substantially slower, even at elevated temperatures, which limits performance of annealing replica-exchange methods.^8^ Increased entropic barriers reduce the rate of interconversion between states within the temperature ladder, leading to markedly longer convergence times. Under these conditions, importance sampling techniques may outperform T-REMD in reaching equilibrium.

The high accuracy of T-REMD observed here also stems from the fact that its primary observable is the folded fraction, which yields precise thermodynamic estimates when the melting temperature lies within the temperature ladder, typically spanning ~270 K to 400–500 K, or close to it. This corresponds to systems in which the folded-state population spans a reasonable dynamic range, approximately 10–90 %. For systems (and FFs), where the folding free energy significantly deviates from 0 kcal/mol for all simulated temperature, thermodynamic estimates based on folded fractions become unreliable. In such cases, importance sampling methods, which directly estimate the ΔG°_fold_ rather than relying on folded populations, offer a clear advantage. They can therefore provide meaningful benchmarks even for challenging systems such as the UNCG TL, whose stability is often underestimated by current RNA FFs, resulting in positive ΔG°_fold_ and negligible folded populations within accessible temperature ranges.^12^

Our results emphasize that benchmarking RNA FFs must pair each test system with a sampling method capable of achieving quantitative convergence. For the GAGA TL, T-REMD simulations on the tens-of-microseconds timescale yielded a converged ensemble at the reference temperature of 298 K. From this ensemble, the folded fraction and ΔG°_fold_ were estimated using an εRMSD-based criterion to define the native state. Although this structural definition is reasonable, rigorous comparison with experimental data requires verifying that the simulated folded (i.e., native-like) ensemble corresponds to the experimentally observed state. Experimental melting temperatures and ΔG°_fold_ values – or, in the case of the 8-mer GAGA TL, their estimates based on Turner’s parameters – are typically derived from UV–vis melting experiments that monitor the hyperchromicity effect at 260 nm. For the 8-mer GAGA TL, Turner-based estimates indicate ΔG°_fold_ of –0.9 ± 0.2 kcal/mol at 300 K,^16, 54–57^ a native-state population of 82 ± 4 %, and a melting temperature of 311.8 K, providing a widely accepted indirect thermodynamic reference derived from UV–vis melting data. Achieving a truly quantitative comparison would therefore require evaluating the spectral response of individual conformational clusters populated in the simulations (see Supported Information for details on misfolded states). Future work will focus on bridging this gap by predicting UV–vis absorption spectra for representative states of the 298 K ensemble using TD-DFT calculations.

## Supporting information

Supporting Information to the article

## ASSOCIATED CONTENT

### Supporting Information

The Supporting Information is available free of charge via the Internet at http://pubs.acs.org/ and containing Supporting Tables and Figures.

## AUTHOR INFORMATION

## ACKNOWLEDGMENT

This work was supported by the Czech Science Foundation to V.M. and J.S. (grant number 23-05639S). This research also received the support of EXA4MIND, a European Union’s Horizon Europe Research and Innovation programme under grant agreement N° 101092944 (V.M., M.O. and P.B). Views and opinions expressed are however those of the author(s) only and do not necessarily reflect those of the European Union or the European Commission. Neither the European Union nor the granting authority can be held responsible for them. P.K., M.O. and P.B. were also supported by ERDF/ESF project TECHSCALE (No. CZ.02.01.01/00/22_008/0004587).

