## Supporting Information to the article for "Toward Accurate RNA Folding Thermodynamics: Evaluation of Enhanced Sampling Methods for Force Field Benchmarking"

SUPPORTING TABLES

**Table S1:** Temperature-dependent populations of states in the T-REMD simulations initiated from the folded ensemble from final 5 μs. For each temperature, the table reports the percentage of frames in the native conformations ("Native"), in conformations with a four-bases stacked loop ("4-stack"), in cross-stem loop conformations ("Cross"), in non-native loop conformations ("Loop"), in single-stranded conformations ("Ss"), and in frames that could not be reliably assigned ("Unassigned"). State definitions are based on εRMSD clustering with the native state defined relative to the experimental hairpin structure (seem Methods in the main text).

| Temperature [K] | Native | 4-stack | Cross | Loop | Ss | Unassigned |
|---|---|---|---|---|---|---|
| 278.0 | 27.0 | 15.9 | 3.6 | 15.0 | 12.0 | 26.5 |
| 279.9 | 26.5 | 15.3 | 3.7 | 15.2 | 11.9 | 27.5 |
| 281.9 | 26.3 | 14.7 | 3.5 | 15.2 | 11.5 | 28.8 |
| 284.0 | 26.0 | 14.0 | 3.3 | 15.5 | 11.3 | 29.9 |
| 286.1 | 26.1 | 13.4 | 3.2 | 15.5 | 11.1 | 30.7 |
| 288.3 | 25.4 | 12.9 | 3.1 | 15.6 | 10.9 | 32.2 |
| 290.4 | 24.8 | 12.5 | 3.0 | 15.7 | 10.7 | 33.3 |
| 292.5 | 24.0 | 11.8 | 2.8 | 15.9 | 10.5 | 35.0 |
| 294.7 | 23.5 | 11.5 | 2.8 | 15.8 | 10.3 | 36.3 |
| 296.8 | 23.2 | 10.9 | 2.6 | 15.6 | 10.2 | 37.5 |
| 299.0 | 22.5 | 10.5 | 2.5 | 15.5 | 10.3 | 38.7 |
| 301.1 | 22.0 | 10.1 | 2.4 | 15.7 | 10.2 | 39.6 |
| 303.3 | 21.3 | 9.6 | 2.2 | 15.7 | 10.2 | 41.0 |
| 305.5 | 20.7 | 9.1 | 2.1 | 15.6 | 9.9 | 42.6 |
| 307.7 | 19.9 | 8.6 | 2.0 | 15.5 | 9.9 | 44.0 |
| 310.0 | 19.2 | 8.3 | 1.9 | 15.5 | 9.8 | 45.3 |
| 312.3 | 18.5 | 7.6 | 1.8 | 15.5 | 9.5 | 47.0 |
| 314.6 | 18.0 | 7.3 | 1.7 | 15.3 | 9.4 | 48.2 |
| 317.0 | 17.5 | 6.8 | 1.6 | 15.1 | 9.2 | 49.8 |
| 319.3 | 16.5 | 6.6 | 1.5 | 14.6 | 9.0 | 51.8 |
| 321.9 | 15.6 | 6.1 | 1.3 | 14.3 | 8.9 | 53.8 |
| 324.4 | 14.9 | 5.7 | 1.3 | 13.9 | 8.7 | 55.5 |
| 326.8 | 14.1 | 5.3 | 1.1 | 13.5 | 8.7 | 57.3 |
| 329.3 | 13.4 | 5.1 | 1.1 | 12.7 | 8.4 | 59.4 |
| 331.9 | 12.4 | 4.9 | 0.9 | 12.5 | 8.2 | 61.1 |
| 334.4 | 11.7 | 4.4 | 0.9 | 12.2 | 8.0 | 62.8 |
| 337.0 | 11.1 | 4.1 | 0.8 | 11.8 | 7.8 | 64.4 |
| 339.6 | 10.3 | 3.8 | 0.7 | 11.2 | 7.7 | 66.3 |
| 342.2 | 9.5 | 3.4 | 0.6 | 10.7 | 7.4 | 68.4 |
| 344.9 | 8.8 | 3.2 | 0.5 | 10.4 | 7.2 | 69.9 |
| 347.6 | 8.0 | 2.9 | 0.5 | 9.9 | 7.0 | 71.7 |
| 350.3 | 7.3 | 2.7 | 0.4 | 9.5 | 6.8 | 73.4 |
| 353.1 | 6.5 | 2.6 | 0.4 | 9.1 | 6.6 | 74.8 |

| | | | | | | |
|---|---|---|---|---|---|---|
| 355.9 | 5.9 | 2.4 | 0.4 | 8.5 | 6.4 | 76.4 |
| 358.7 | 5.4 | 2.2 | 0.3 | 8.0 | 6.2 | 78.0 |
| 361.6 | 4.9 | 2.1 | 0.2 | 7.5 | 5.8 | 79.6 |
| 364.5 | 4.3 | 1.9 | 0.2 | 6.9 | 5.5 | 81.2 |
| 367.4 | 3.9 | 1.8 | 0.2 | 6.5 | 5.2 | 82.4 |
| 370.4 | 3.5 | 1.5 | 0.2 | 5.9 | 4.9 | 84.0 |
| 373.4 | 3.0 | 1.3 | 0.1 | 5.6 | 4.6 | 85.4 |
| 376.4 | 2.6 | 1.2 | 0.1 | 5.2 | 4.4 | 86.5 |
| 379.7 | 2.4 | 1.1 | 0.1 | 4.6 | 4.1 | 87.6 |
| 382.9 | 2.0 | 1.1 | 0.1 | 4.3 | 3.8 | 88.7 |
| 386.1 | 1.7 | 1.0 | 0.1 | 3.8 | 3.5 | 89.9 |
| 389.3 | 1.4 | 0.9 | 0.1 | 3.5 | 3.3 | 90.8 |
| 392.6 | 1.3 | 0.8 | 0.1 | 3.3 | 3.0 | 91.5 |
| 396.2 | 1.1 | 0.7 | 0.1 | 3.0 | 2.7 | 92.4 |
| 399.6 | 1.0 | 0.6 | 0.1 | 2.7 | 2.5 | 93.1 |
| 403.1 | 0.8 | 0.5 | 0.1 | 2.4 | 2.2 | 93.9 |
| 406.6 | 0.7 | 0.5 | 0.1 | 2.2 | 2.1 | 94.4 |
| 410.1 | 0.6 | 0.4 | 0.0 | 1.9 | 1.9 | 95.0 |
| 413.7 | 0.5 | 0.4 | 0.0 | 1.6 | 1.8 | 95.6 |
| 417.3 | 0.5 | 0.3 | 0.0 | 1.4 | 1.6 | 96.2 |
| 421.0 | 0.4 | 0.3 | 0.0 | 1.2 | 1.6 | 96.5 |
| 424.9 | 0.3 | 0.3 | 0.0 | 1.1 | 1.5 | 96.8 |
| 428.7 | 0.3 | 0.3 | 0.0 | 0.9 | 1.4 | 97.1 |
| 432.5 | 0.3 | 0.3 | 0.0 | 0.8 | 1.4 | 97.2 |
| 436.4 | 0.2 | 0.2 | 0.0 | 0.7 | 1.3 | 97.6 |
| 440.4 | 0.2 | 0.2 | 0.0 | 0.6 | 1.2 | 97.8 |
| 444.4 | 0.2 | 0.2 | 0.0 | 0.5 | 1.1 | 98.1 |
| 448.5 | 0.1 | 0.1 | 0.0 | 0.4 | 0.9 | 98.4 |
| 452.6 | 0.1 | 0.1 | 0.0 | 0.4 | 0.9 | 98.4 |
| 456.8 | 0.1 | 0.1 | 0.0 | 0.4 | 0.8 | 98.6 |
| 461.0 | 0.0 | 0.1 | 0.0 | 0.3 | 0.7 | 98.8 |

**Table S2:** Temperature-dependent populations of states in the T-REMD simulations initiated from the unfolded ensemble from final 5 μs. See Table S1 for more details about state definitions.

| Temperature [K] | Native | 4-stack | Cross | Loop | Ss | Unassigned |
|---|---|---|---|---|---|---|
| 278.0 | 25.4 | 14.9 | 9.3 | 14.3 | 10.5 | 25.5 |
| 279.9 | 25.5 | 14.2 | 9.1 | 14.3 | 10.4 | 26.6 |
| 281.9 | 25.1 | 13.7 | 8.6 | 14.5 | 10.4 | 27.8 |
| 284.0 | 24.4 | 13.2 | 8.5 | 14.5 | 10.2 | 29.2 |
| 286.1 | 24.0 | 12.8 | 8.1 | 14.6 | 10.1 | 30.4 |
| 288.3 | 23.5 | 12.1 | 7.7 | 15.1 | 9.9 | 31.6 |
| 290.4 | 23.2 | 11.5 | 7.5 | 15.2 | 9.9 | 32.8 |
| 292.5 | 23.0 | 11.0 | 7.1 | 15.2 | 9.7 | 34.1 |
| 294.7 | 21.9 | 10.6 | 7.0 | 15.5 | 9.6 | 35.3 |
| 296.8 | 21.5 | 10.0 | 7.1 | 15.6 | 9.5 | 36.3 |
| 299.0 | 21.3 | 9.4 | 6.7 | 15.6 | 9.4 | 37.5 |
| 301.1 | 20.4 | 8.9 | 6.5 | 15.5 | 9.4 | 39.2 |
| 303.3 | 20.2 | 8.1 | 6.3 | 15.4 | 9.6 | 40.5 |
| 305.5 | 19.7 | 7.8 | 6.0 | 15.6 | 9.6 | 41.4 |
| 307.7 | 18.9 | 7.5 | 5.6 | 15.4 | 9.5 | 43.1 |
| 310.0 | 18.2 | 7.1 | 5.3 | 15.4 | 9.1 | 44.9 |
| 312.3 | 17.6 | 6.6 | 4.9 | 15.5 | 9.1 | 46.2 |
| 314.6 | 16.9 | 6.1 | 4.5 | 15.2 | 8.8 | 48.5 |
| 317.0 | 16.1 | 5.8 | 4.2 | 15.1 | 8.7 | 50.1 |
| 319.3 | 15.5 | 5.5 | 3.9 | 14.7 | 8.5 | 51.8 |
| 321.9 | 14.8 | 5.2 | 3.6 | 14.4 | 8.3 | 53.8 |
| 324.4 | 13.9 | 4.8 | 3.4 | 14.1 | 8.2 | 55.6 |
| 326.8 | 12.9 | 4.5 | 3.2 | 13.6 | 8.1 | 57.7 |
| 329.3 | 12.1 | 4.2 | 2.9 | 13.3 | 7.9 | 59.6 |
| 331.9 | 11.5 | 3.9 | 2.7 | 13.1 | 7.8 | 61.0 |
| 334.4 | 10.7 | 3.8 | 2.5 | 12.8 | 7.7 | 62.6 |
| 337.0 | 10.0 | 3.6 | 2.4 | 12.3 | 7.6 | 64.1 |
| 339.6 | 9.4 | 3.3 | 2.2 | 12.0 | 7.4 | 65.8 |
| 342.2 | 8.9 | 2.9 | 1.9 | 11.4 | 7.1 | 67.8 |
| 344.9 | 8.3 | 2.7 | 1.9 | 10.8 | 6.8 | 69.5 |
| 347.6 | 7.7 | 2.5 | 1.7 | 10.2 | 6.5 | 71.3 |
| 350.3 | 6.9 | 2.3 | 1.6 | 9.7 | 6.3 | 73.2 |
| 353.1 | 6.2 | 2.1 | 1.4 | 9.0 | 6.2 | 75.1 |
| 355.9 | 5.6 | 1.9 | 1.3 | 8.4 | 6.1 | 76.7 |
| 358.7 | 5.2 | 1.6 | 1.2 | 8.0 | 5.7 | 78.3 |
| 361.6 | 4.8 | 1.5 | 1.1 | 7.4 | 5.5 | 79.8 |
| 364.5 | 4.3 | 1.3 | 0.9 | 6.9 | 5.4 | 81.2 |
| 367.4 | 4.0 | 1.2 | 0.8 | 6.3 | 5.2 | 82.6 |
| 370.4 | 3.7 | 1.1 | 0.7 | 5.8 | 5.0 | 83.7 |

| | | | | | | |
|---|---|---|---|---|---|---|
| 373.4 | 3.3 | 1.0 | 0.6 | 5.5 | 4.7 | 85.0 |
| 376.4 | 3.0 | 0.9 | 0.5 | 4.9 | 4.6 | 86.1 |
| 379.7 | 2.6 | 0.9 | 0.5 | 4.4 | 4.5 | 87.1 |
| 382.9 | 2.3 | 0.8 | 0.5 | 3.9 | 4.3 | 88.3 |
| 386.1 | 2.0 | 0.8 | 0.4 | 3.5 | 4.0 | 89.3 |
| 389.3 | 1.7 | 0.7 | 0.3 | 3.2 | 3.7 | 90.4 |
| 392.6 | 1.3 | 0.7 | 0.2 | 2.9 | 3.6 | 91.3 |
| 396.2 | 1.1 | 0.6 | 0.2 | 2.5 | 3.3 | 92.2 |
| 399.6 | 0.9 | 0.5 | 0.2 | 2.2 | 3.2 | 93.0 |
| 403.1 | 0.8 | 0.5 | 0.2 | 1.9 | 2.9 | 93.7 |
| 406.6 | 0.7 | 0.4 | 0.1 | 1.6 | 2.7 | 94.4 |
| 410.1 | 0.6 | 0.4 | 0.1 | 1.3 | 2.4 | 95.1 |
| 413.7 | 0.5 | 0.4 | 0.1 | 1.2 | 2.2 | 95.6 |
| 417.3 | 0.5 | 0.4 | 0.1 | 1.0 | 2.0 | 96.1 |
| 421.0 | 0.4 | 0.3 | 0.1 | 0.8 | 1.8 | 96.4 |
| 424.9 | 0.4 | 0.3 | 0.1 | 0.8 | 1.7 | 96.8 |
| 428.7 | 0.3 | 0.3 | 0.1 | 0.6 | 1.5 | 97.2 |
| 432.5 | 0.2 | 0.2 | 0.0 | 0.5 | 1.3 | 97.7 |
| 436.4 | 0.2 | 0.1 | 0.0 | 0.5 | 1.2 | 97.9 |
| 440.4 | 0.2 | 0.2 | 0.0 | 0.4 | 1.2 | 98.1 |
| 444.4 | 0.1 | 0.1 | 0.0 | 0.4 | 1.1 | 98.2 |
| 448.5 | 0.1 | 0.1 | 0.0 | 0.3 | 1.0 | 98.4 |
| 452.6 | 0.1 | 0.1 | 0.0 | 0.3 | 0.9 | 98.6 |
| 456.8 | 0.1 | 0.1 | 0.0 | 0.2 | 0.9 | 98.7 |
| 461.0 | 0.0 | 0.1 | 0.0 | 0.2 | 0.8 | 98.9 |

SUPPORTING FIGURES

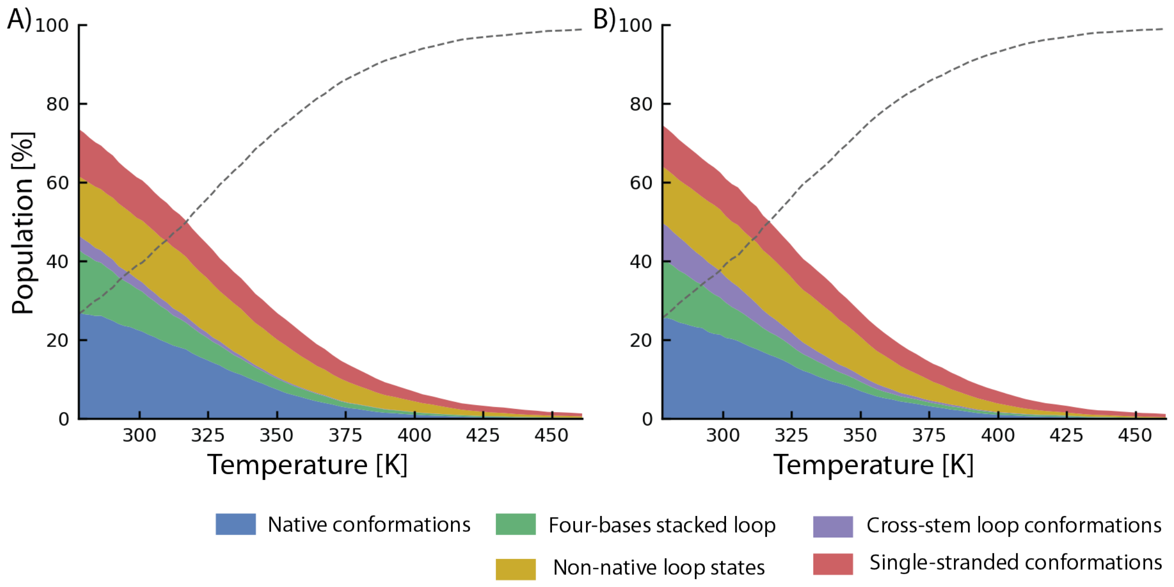

**Figure S1:** Temperature-dependent populations of conformations in the T-REMD simulations initiated from the folded (A) and unfolded ensemble (B) from final 5 µs. The dashed gray line corresponds to Unassigned states, i.e., frames that cannot be reliably assigned by the clustering approach (algorithm introduced by Rodriguez and Laio[1] in combination with the εRMSD metric[2]; see ref [3] for more details). Population of all states are summed to 100%. Numerical values for all states and temperatures are reported in Tables S1 and S2. State definitions are based on εRMSD clustering with the native state defined relative to the experimental hairpin structure (see Methods in the main text for details).

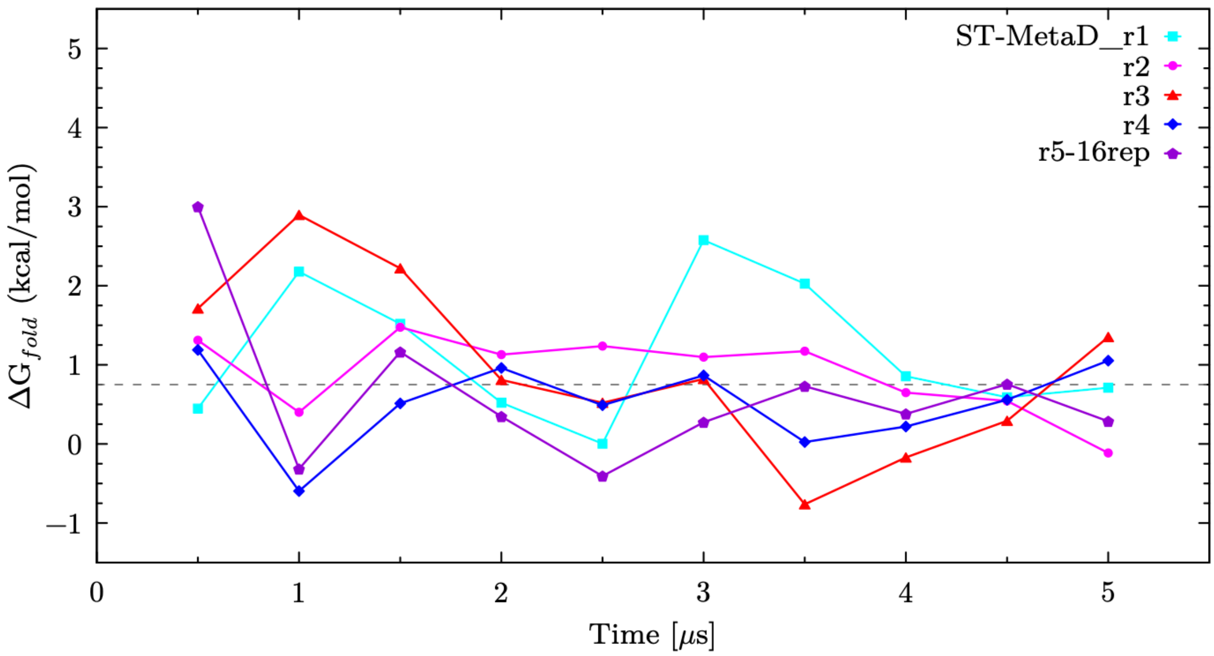

**Figure S2:** Time evolution of folding free energies (ΔG°$_{fold}$) from five 5 μs-long ST-MetaD simulations estimated from the corresponding reweighted ensembles using standard εRMSD (see Methods in the main text), with each line representing values computed using the instantaneous bias accumulated up to the given time.

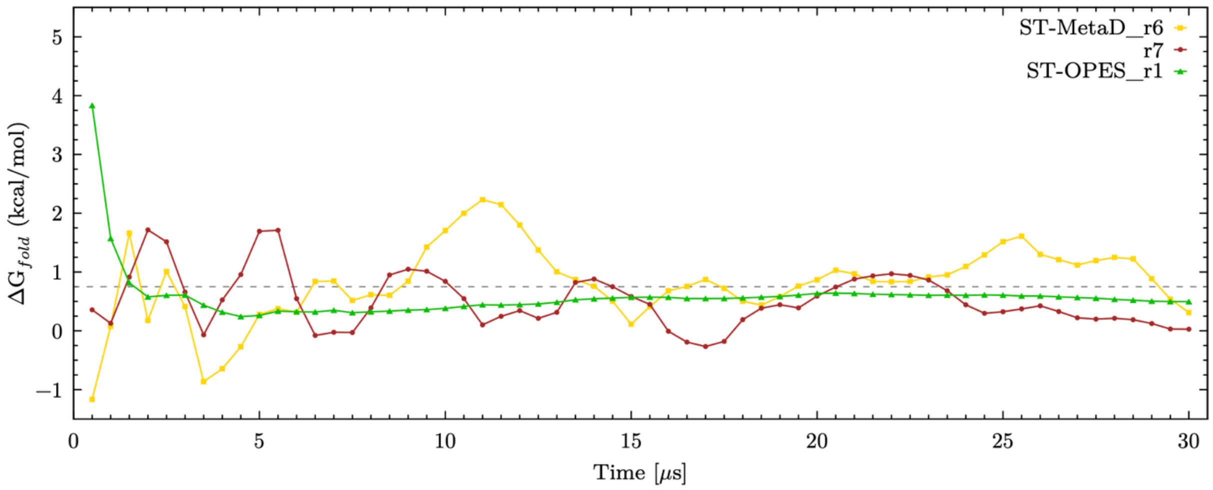

**Figure S3:** Time evolution of folding free energies (ΔG°$_{fold}$) from two 30 μs-long ST-MetaD simulations and one 30 μs-long ST-OPES simulation estimated from the corresponding reweighted ensembles using standard εRMSD (see Methods in the main text), with each line representing values computed using the instantaneous bias accumulated up to the given time.
